# Inflammasome Profiling in Human Natural Killer Cells under Pro-inflammatory Stimuli and Solid Organ Transplantation

**DOI:** 10.1101/2024.04.17.589865

**Authors:** Antonio Astorga-Gamaza, Inés Muela-Zarzuela, Juan Miguel Suárez-Rivero, Alexis Varin, Maarten Naesens, Alessandra Tammaro, Jesper Kers, Juan López-Pérez, Raquel de la Varga-Martínez, Auxiliadora Mazuecos, Baptiste Lamarthée, Mario D. Cordero

## Abstract

Innate immunity relies on inflammasomes as key mediators of host defense, orchestrating the release of pro-inflammatory cytokines and triggering pyroptotic cell death in response to harmful stimuli. Although inflammasome activity has been extensively studied in myeloid cells, its role in natural killer (NK) cells remains underexplored. This study demonstrates that human primary NK cells can functionally activate inflammasomes both in vitro and in vivo, including in patients undergoing organ transplantation. Ex vivo stimulation with nigericin and the dipeptidyl peptidases (DPP) inhibitor Talabostat (Val-boroPro) induces pyroptotic cell death in a subset of NK cells. This is marked by the cleavage and activation of gasdermin D, a lytic pore-forming protein essential for pyroptosis. Accompanying gasdermin D activation, significant levels of lactate dehydrogenase (LDH) and residual amounts of interleukin-18 (IL-18) are released. The detection of activated caspase-4 further indicates that these processes are mediated through non-canonical inflammasome pathways in NK cells. Notably, CD56^dim^ and CD56^bright^ NK cell subsets exhibit distinct responses to pro-inflammatory stimulation. In patients with renal dysfunction, sustained inflammasome activation, particularly involving NLRP1 and NLRP3, is observed in NK cells, with a shift toward a more pro-inflammatory phenotype following kidney transplantation. Single-cell RNA sequencing analyses further reveal persistently elevated expression of caspase-4 and gasdermin-D in transplant recipients experiencing rejection and microvascular inflammation. These findings highlight the underappreciated role of NK cells in inflammasome-driven inflammation, underscoring their importance in both basic research and clinical contexts.

## Introduction

Inflammation is part of the body’s defense response to harmful stimuli. However, exacerbated inflammation is also associated with a range of diseases. In this regard, the activity of natural killer (NK) cells has been shown to highly depend on the context, helping to reduce or even promote inflammatory processes [1].

NK cells are innate lymphoid cells characterized by a huge diversity of phenotypes and functions [2, 3], although they are primarily categorized into two main subpopulations based on the neural cell adhesion molecule CD56 density. CD56^dim^ NK cells are known for their potent cytolytic activities, and CD56^bright^ NK cells are distinguished by their cytokine secretion capabilities [4]. They play a pivotal role in immune surveillance, effectively targeting malignant and infected cells to control intracellular pathogens and shaping adaptive immune responses [5]. Equipped with a huge array of cell surface receptors, NK cells sense stress signals in target cells, distinguishing between ‘self’ and ‘non-self’ or ‘aberrant’ targets under normal physiological conditions [6]. Upon activation, NK cells release cytotoxic granules containing perforin and granzyme. Also, they secrete interferon (IFN)-γ and tumor necrosis factor (TNF), and immunoregulatory molecules such as interleukin (IL)-10 and the growth factor GM-CSF, as well as chemokines including CCL3 (MIP-1α), CCL4 (MIP-1β), CCL5 (RANTES), and CXCL8 (IL-8) [7]. Through these mechanisms, NK cells can drive inflammation and modulate the activity of other immune cells.

A key part of the inflammatory process and the innate immune response is the assembly of inflammasomes. These cytosolic multi-protein complexes regulate tissue inflammation in response to multiple cellular stress patterns, including pathogen-associated molecular patterns (PAMPs) such as bacterial lipopolysaccharide (LPS), or cellular damage-associated molecular patterns (DAMPs), which include extracellular ATP and changes in intracellular ion concentrations. In this regard, inflammasome biology has been mostly characterized in macrophages and at least partially studied in other cell types such as endothelial cells, fibroblasts, epithelial cells, and T cells [8–11]. However, it is unknown whether NK cells are equipped with functional inflammasome components, and thus, this could represent an underexplored arm of the diverse immune mechanisms mediated by these cells and an important source of inflammation.

The first steps in the triggering of inflammasomes comprise the detection of the damage patterns via sensors of several families such as the NOD-like (NLR), Toll-like (TLR), AIM2-like (ALR), or the caspase recruitment domain (CARD)-containing family of proteins, that include the members NLRP1, NLRP3, NLRC4, AIM2, and CARD8, among others [12]. Following activation, some sensors interact with adaptor proteins such as ASC (apoptosis-associated speck-like protein containing a CARD), composing a pre-complex that then recruits inflammatory caspase-1. Next to terminal assembly, the activation of caspase-1 in the inflammasome can lead to the proteolytic cleavage and maturation of the IL-1 family cytokines IL-1β and IL-18 [13]. This pathway is known as the “canonical” activation. Another usual key event after the inflammasome activation is the cleavage of gasdermins, a family of pore-forming proteins, into its N-terminal active domain, which induces a proinflammatory form of regulated cell death termed pyroptosis [14]. Inflammasome activation can also occur in a named “non-canonical” way, with caspases-4 and −5 in humans directly promoting pyroptosis [15, 16].

Herein, we conducted a transcriptomic analysis of NK cells derived from various tissues, revealing the expression of genes encoding several components of the inflammasome. Additionally, we provide proteomic evidence demonstrating the expression of all the components for the NLRP3 and NLRP1 inflammasome in both CD56^dim^ and CD56^bright^ NK cell subsets. More importantly, we demonstrate the functionality of these inflammasomes in NK cells, mainly evidenced by the induction of pyroptotic cell death and activation of gasdermin-D. Interestingly, we observe distinct responses to pro-inflammatory stimuli between the two NK cell subpopulations. Furthermore, we identify signs of inflammasome activation in NK cells from patients with renal dysfunction, which persist despite immunosuppression after kidney transplantation. Our findings suggest that NK cells should be considered in inflammation research and as potential therapeutic targets, as they might play a role in both pathogen defense and inflammatory disorders through inflammasome signaling and pyroptosis.

## RESULTS

### Expression of inflammation-related genes in human NK cells from different tissues

To gain insight into the potential of human NK cells in shaping inflammatory responses and constituting functional inflammasomes, we conducted a transcriptomic analysis of NK cells from diverse tissues using a publicly available RNA-seq dataset [3]. Cells were isolated from donors without cancer or any other chronic disease, and seronegative for hepatitis B, C, and HIV viral infections [3]. We assessed the expression of a list of genes related to inflammation, considering a reference list [17]. From a total of 2,686 genes, we found expression of 1,619 in NK cells from blood, bone marrow, lymph nodes, lungs, and spleen. These genes appeared to be involved in different processes such as the regulation of the NF-κB activity, apoptosis, TNF signaling, cellular response to lipopolysaccharide, NOD-like and Toll-like receptor signaling, IL-6 and IL-1β production (adjusted p value < 0.05) (**Figure 1A**). Next, we performed a detailed analysis of the expression of short-listed genes reported to play a role in activating inflammasome multiprotein complexes at different stages (**Figure 1B, C, D, E, and F**). Interestingly, we noticed the expression in NK cells of guanylate-binding proteins (GBP), which target intracellular pathogens and can mediate host defense via inflammasome induction [18, 19]. Moreover, we found expression of other receptors of DAMPs and PAMPs such as *TLR-2-3* and *4*, *NLRP1*, *NLRP3*, *NLRP6*, *NLRC3*, and *NLRC4,* the interferon-inducible protein *AIM-2* and *CARD*. NK cells also expressed *PYCARD*, the gene coding for ASC protein, required to constitute canonical inflammasomes, and the serin proteases *DPP8* and *DPP9*, whose pharmacological inhibition by Val-boroPro (Vbp) (Talabostat) is reported to activate both NLRP1 and CARD8 inflammasomes [20, 21]. Furthermore, NK cells from all analyzed tissues expressed downstream effectors in the inflammasome pathway such as *CASPASE-1*, *IL-1β*, *IL-18*, and *GASDERMIN D* genes (**Figure 1B, C, D, E, and F**). Some significant differences were noticed between the NK subpopulations CD56^dim^ and CD56^bright^. This was the case for *GBP5* and *TLR3* in the blood (**Figure 1B**), and *CASPASE-1* in the lung (**Figure 1C**), with higher expression in CD56^dim^ NK cells. However, larger datasets are needed to confirm and reveal differences between different NK subpopulations as NK cells exhibit an intrinsic, highly heterogeneous nature intra- and inter-individually [2].

**Figure 1.**
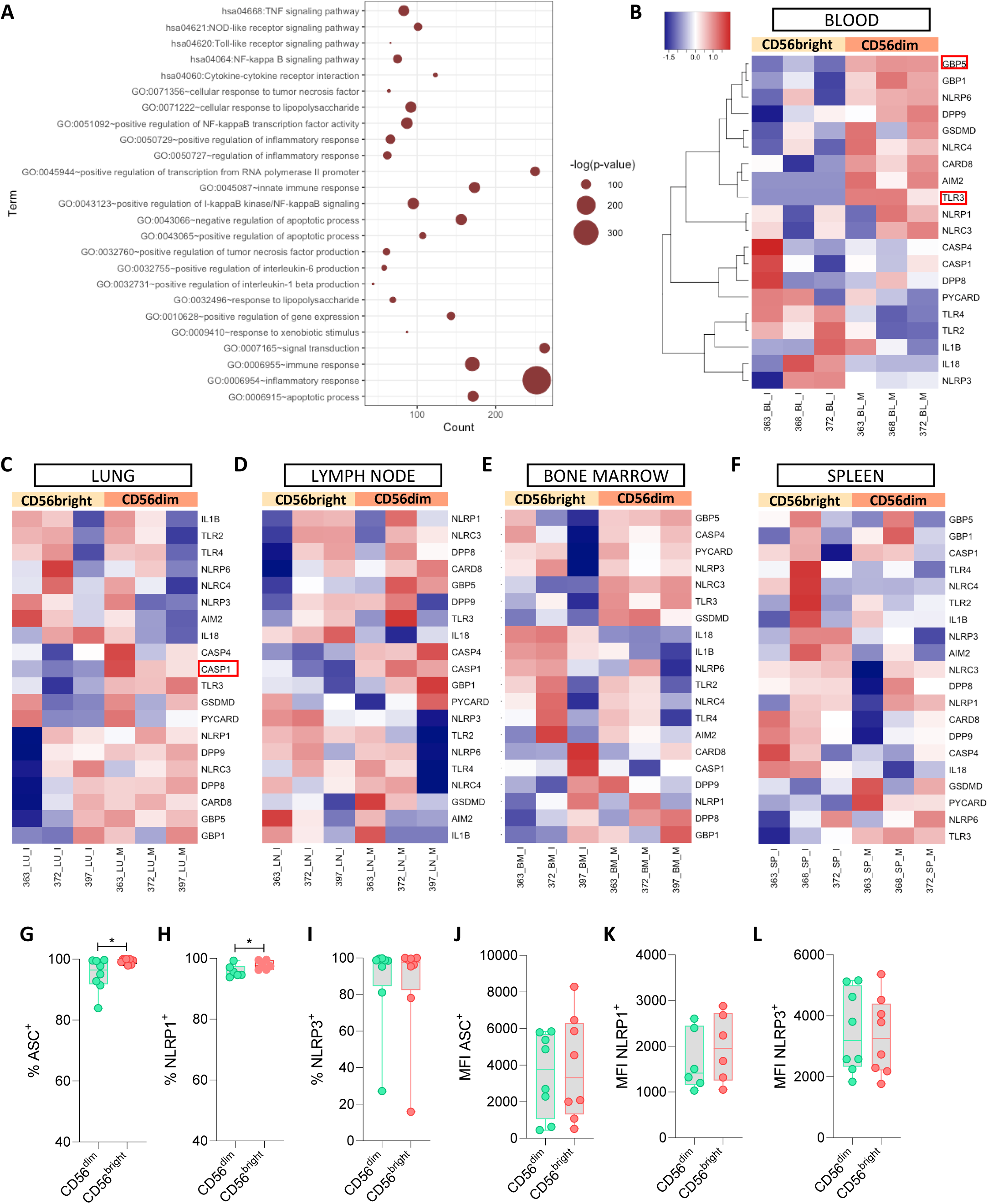
RNA-Seq analysis of inflammatory transcriptional signatures in circulating and tissue-resident human NK cells, and protein expression of the NLRP1 and NLRP3 inflammasome sensors. Cells were isolated from donors without cancer or any other chronic disease, and seronegative for hepatitis B, C, and HIV viral infections. **A)** Significance of the most represented immune pathways according to the expression of a list of 2,686 inflammation-related genes. Functional gene annotation was performed using DAVID. **B)** Heatmap of the expression of short-listed genes key in the activation and triggering of effector functions of inflammasomes in circulating NK cells. An analysis of the differential expression of these genes between CD56^bright^CD16^−^ and CD56^dim^CD16^+^ NK cell subsets is also shown, and genes showing a different expression pattern between both NK subpopulations are marked with a red box. The same is shown for **C)** NK cells in the lung, D) NK cells in the lymph nodes, **E)** NK cells in the bone marrow, and **F)** NK cells in the spleen. Differentially expressed genes were identified using linear models (Limma-Voom) and P values adjusted for multiple comparisons by applying the Benjamini-Hochberg correction. **G)** Frequency (%) of expression of ASC in sorted CD56^dim^CD16^+^ and CD56^bright^CD16^-/dim^ NK cells (n=8), **H)** Frequency of NLRP1, and **I)** Frequency of NLRP3. Mean Fluorescence Intensity (MFI) signal for each marker in **J)**, **K)**, and **L)**. Median with range is represented. Statistical comparisons were performed using the Wilcoxon matched-pairs signed-rank test.

Next, we aimed to assess the expression of the protein sensors NLRP1 and NLRP3, which have been implicated in the regulation and pathogenesis of many conditions. We studied the expression of the proteins NLRP1, NLRP3, and the adaptor ASC in NK cells by flow cytometry, using blood-derived primary NK cells from healthy donors previously isolated by FACS (fluorescence-activated cell sorting) (**Supplementary Figure 1A**). The median purity of the cells was 90.7% and 94.8% for CD56^dim^ and CD56^bright^ NK cells, respectively. A representative example of staining can be found in **Supplementary Figure 1B-C**. These proteins were widely expressed in circulating NK cells, with slightly higher frequencies of ASC and NLRP1 in the CD56^bright^ subpopulation (**Figure 1G and H**), but no differences in the percentage of NLRP3 (**Figure 1I**) nor in the intensity of expression (mean fluorescence intensity, MFI) of any of them (**Figure 1J, K, and L**). Our results indicate that both the two main CD56^dim^ and CD56^bright^ subpopulations of NK cells express the components needed to constitute functional inflammasomes, and particularly NLRP3 and NLRP1 inflammasomes, suggesting that they may play a role in NK cell-mediated immunity.

### NK cell pyroptosis triggered by specific NLRP1 and NLRP3 inflammasome activators

As a next step, we directly tested the functionality of inflammasomes in NK cells. For that, we treated blood-derived NK cells from healthy volunteers, previously isolated by FACS, with pharmacological activators of the NLRP3 and NLRP1 inflammasomes ex vivo. We included conditions of cells cultured with lipopolysaccharide (LPS) plus ATP (LPS-ATP), nigericin (Nig), and the DPP8/DPP9 inhibitor Vbp for both CD56^dim^ and CD56^bright^ NK subpopulations separately. Cytokine secretion was barely detected, with minimal amounts of IL-18 upon stimulation of CD56^dim^ cells with nigericin (**Figure 2A**) or CD56^bright^ cells with Vbp (**Figure 2B)**. No increment was detected in any of the subpopulations after LPS-ATP stimulation (**Supplementary Figures 2A and B**). IL-1β was also not induced *in vitro*, except for one donor that also showed higher levels of IL-6 (**Supplementary Figures 2C, D, and E**). Besides, we measured other immune mediators by a multiplex Luminex immunoassay. The secretion of IL-10, GM-CSF, and the NK-associated cytotoxic and regulatory molecules IFN-γ, IFN-α, or TNF-α was not detectable in our assay conditions, suggesting that in vitro treatment with nigericin and Vbp does not induce unspecific NK cytotoxicity. Interestingly, ICAM-1, the adhesion receptor and regulator of inflammation, seemed to increase its concentration after LPS-ATP and nigericin treatment of CD56^dim^ cells (**Supplementary Figures 2F and G**). These results suggest that different pro-inflammatory stimuli might differentially affect distinct NK subpopulations.

**Figure 2.**
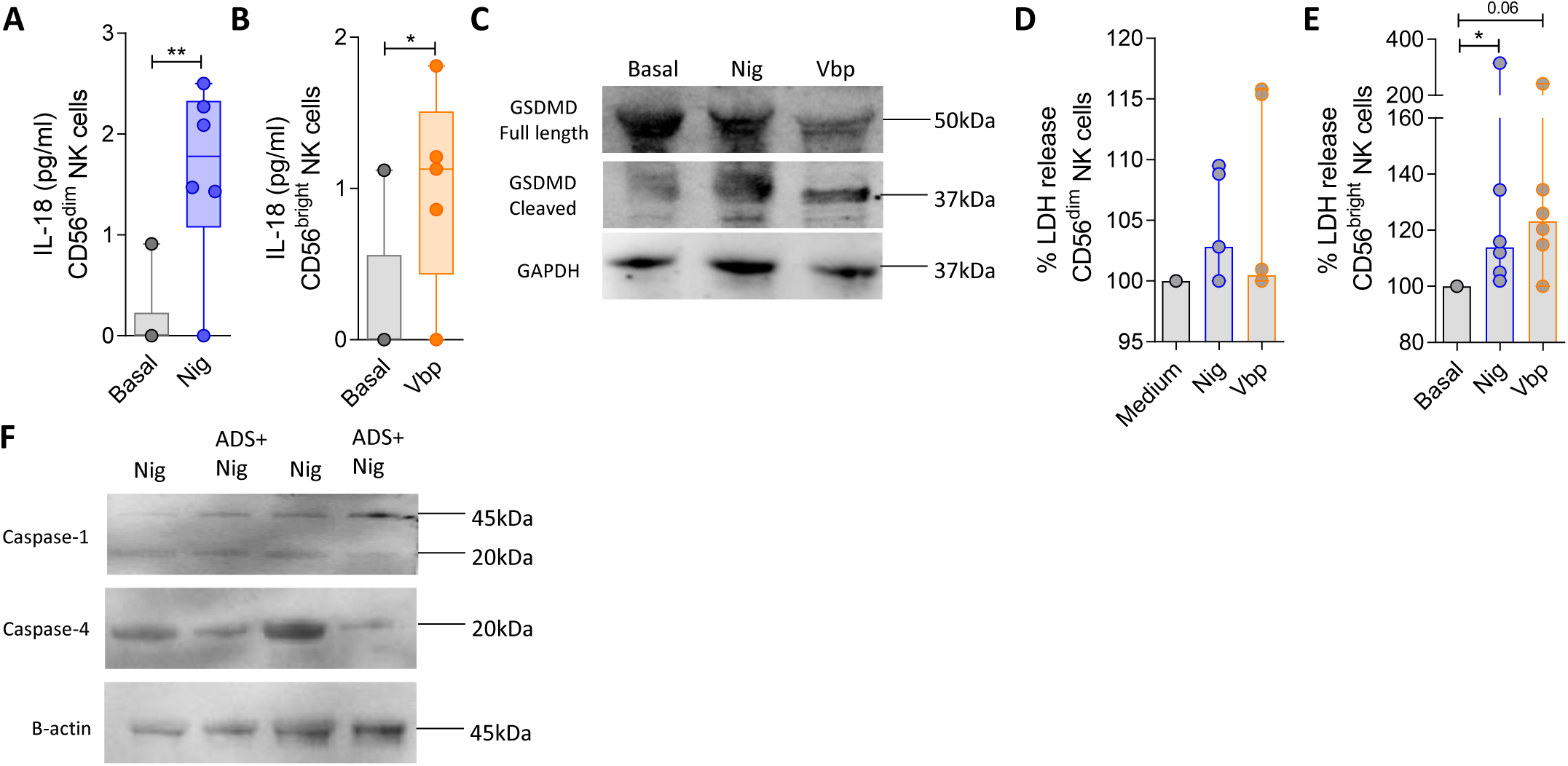
Activation of the NLRP1 and NLRP3 inflammasomes and induction of pyroptosis in human NK cells ex vivo. Isolated NK cells from blood samples were cultured in the presence of the inflammasome activators LPS (1 µg/ml for 4 hours) and ATP (5 mM for 30 mins), nigericin (10 µM for 30 mins) or Vbp (1 µM for 4 hours). After that, the concentration of cytokines of the IL-1 family IL-18 and IL1β as well as lactate dehydrogenase (LDH) release were measured in the culture supernatants. Changes in the cell expression of downstream effectors in the inflammasome cascade such as caspase-1, caspase-4, and gasdermin D were also measured by western blot. **A)** Concentration of IL-18 in cell cultures of untreated CD56^dim^ NK cells or treated with nigericin (n=6). **B)** IL-18 concentration in cell cultures of CD56^bright^ NK cells stimulated with Vbp (n=5). **C)** Western blot analysis of the full length and cleaved form of gasdermin D in CD56^dim^ NK cells in basal conditions and after nigericin and Vbp exposure. Representative result from n=5 independent experiments. **D)** LDH release in cell cultures of stimulated CD56^dim^ NK cells (n=6), and **E)** stimulated CD56^bright^ NK cells (n=6). **F)** Western blot analysis of the active forms of caspase-1 and caspase-4 in NK cells after nigericin exposure and previous treatment with the NLRP1/NLRP3 inflammasome inhibitor ADS032 before the nigericin stimulation.

Next, we assessed cell death by pyroptosis, a key process induced after inflammasome activation. Notably, we detected the activation of the N-terminal fragment of gasdermin D (GSDMD-Cleaved) by western blot after nigericin and Vbp treatment of NK cells (**Figure 2C**). Of note, this was successfully detected only in CD56^dim^ NK cells. However, the release of LDH, which confirms cell death, was more prominent in the CD56^bright^ subpopulation (**Figure 2D and E**), raising the possibility that more aggressive death of CD56^bright^ NK cells may hinder the detection of active gasdermin D. Both pro-inflammatory cytokine secretion and gasdermin-dependent pyroptosis rely on the previous cleavage of inflammatory caspases into their active form [13]. Here, we failed to observe increased active fragments of caspase-1 in NK cells stimulated with nigericin and Vbp in 6 out of 7 experiments. Importantly, in additional experiments we aimed to measure activated caspase-4, and it was successfully detected (**Figure 2F**), suggesting the non-canonical activation of the inflammasomes in NK cells. Moreover, the inhibition of NLRP1 and NLRP3 using the dual inhibitor ADS032 lowered the amount of activated caspase-4 in NK cells (**Figure 2F**).

### NLRP1 and NLRP3 undergo protein reorganization in NK cells under pro-inflammatory stimuli, and their activation does not require ASC oligomerization

To gain further insight into the activation of inflammasomes in NK cells, we assessed the expression of proteins within control and treated cells. The purity of the cells employed in these assays was always higher than 90.0%. Moreover, any remaining cell contaminants in the culture were discarded from our analysis using lineage markers for NK cells in the flow cytometry panel. Treatment of NK cells, CD56^dim^ or CD56^bright^, with LPS-ATP, nigericin, or Vbp, did not alter the frequency of cells expressing NLRP3 (**Figures 3A and B**), NLRP1 (**Figures 3C and D**), or ASC (**Figures 3E and F**). However, changes in the intensity of expression (MFI) were detected in the case of nigericin exposure of CD56^bright^ NK cells, with a reduction in NLRP3 levels (**Figure 3G**). This was not observed for the other proteins or stimuli (**Supplementary Figures 3A, B, C, D, and E**). This may indicate that nigericin and Vbp promote the reorganization of these molecules to assemble the inflammasomes rather than drastically changing their expression levels. Additionally, nigericin may promote a more aggressive pyroptotic death of CD56^bright^ NK cells, which could explain the lower levels of NLRP3 and higher LDH release (as shown in Figure 2D and E). Moreover, we studied the formation of specks by flow cytometry, as previously reported [22]. In the formation of inflammasomes, when canonical activation occurs, ASC interacts with upstream sensors such as NLRP1 and NLRP3, which subsequently trigger its assembly into a structure known as a speck composed of multimers of ASC dimers. The speck can be detected by flow cytometry using the time-of-flight (TOF) signal of ASC, as it correlates with a higher pulse height-to-area (H:A) ratio (**Figure 3H)**. Despite consistent signs of speck in CD56^dim^ NK cells after nigericin stimulation, specking levels were very low (**Figure 3I**). Treatment with Vbp also slightly induced specks in 5 out of 7 independent assays in CD56^bright^ NK cells (**Figure 3J**). We did not detect any sign of speck in LPS-ATP or Vbp-treated CD56^dim^ NK cells (**Supplementary Figures 3F and G**), nor consistently in LPS-ATP and nigericin treated CD56^bright^ subpopulation (**Supplementary Figures 3H and I**). Next, we directly explored and visualized the specking of ASC through fluorescent flow cytometry imaging (Amnis®). Primary monocytes and NK cells treated with nigericin, with or without previous treatment with the NLRP1 and NLRP3 dual inhibitor ADS032, were acquired and gated based on the expression of CD3, CD56, and CD14. As expected, we observed a fraction of monocytes with a cytosolic diffusion pattern of ASC expression (**Figure 3K**), but also a significant fraction showing ASC assembly (**Figure 3L**), which was lower in the presence of the dual inflammasome inhibitor ADS032 (**Figure 3M**). Unlike monocytes, even though we still detected some NK cells with ASC specking (**Figures 3N and O**), ASC assembly was rare in NK cells (**Figure 3P**), confirming the results of our flow cytometry assay.

**Figure 3.**
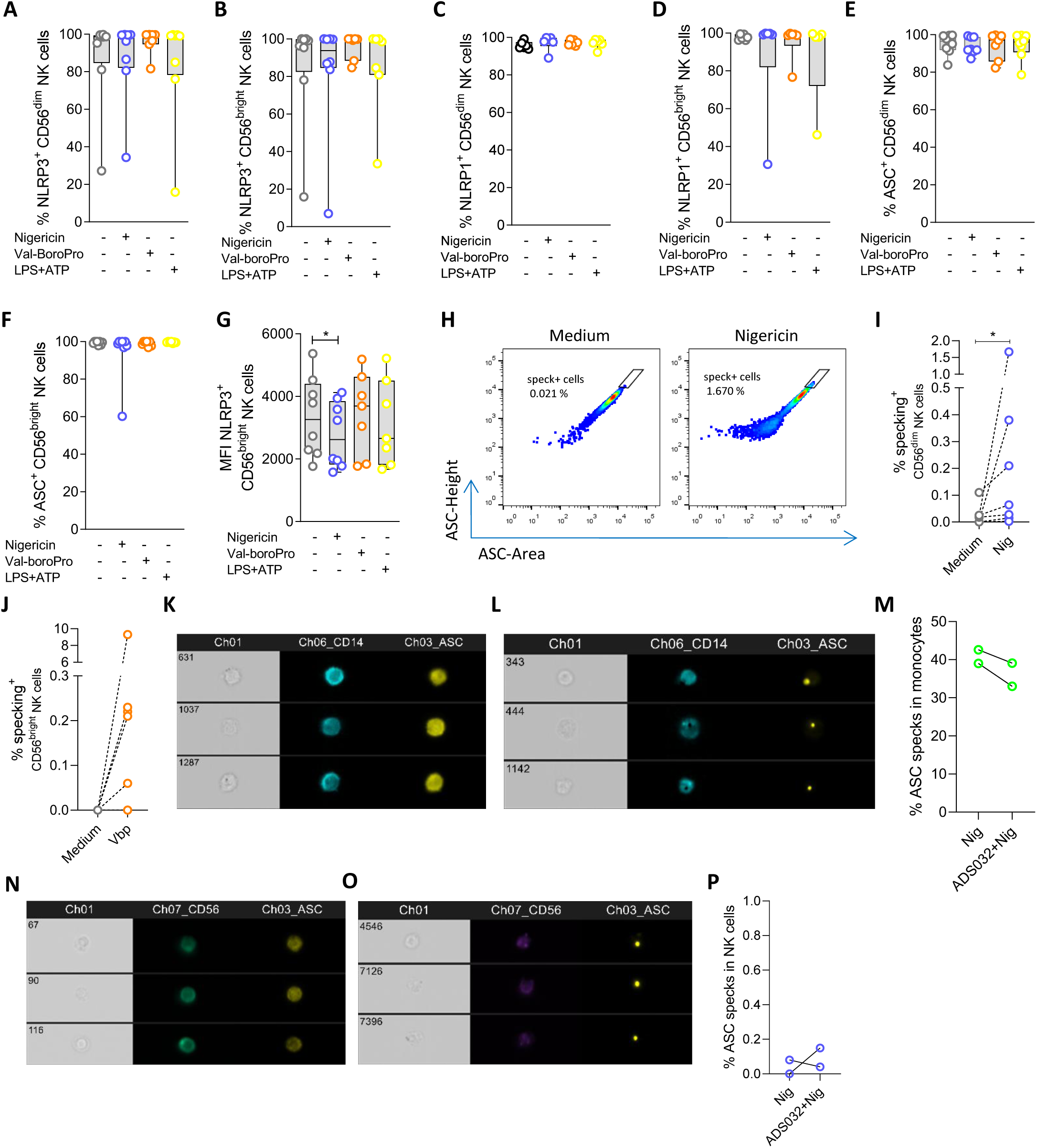
NLRP1, NLRP3, and adaptor ASC inflammasome protein levels in NK cells under different proinflammatory stimuli. **A)** to **G)** represent the frequency and MFI signal for the expression of NLRP1, NLRP3 and ASC in CD56^dim^ and CD56^bright^ NK cells in non-stimulated and under pro-inflammatory stimuli with LPS-ATP, nigericin, and Vbp conditions (n=8), measured by flow cytometry. Graphs represent median with range. Statistical comparisons were performed using the Wilcoxon matched-pairs signed-rank test. **H)** Representative flow cytometry plots of the time-of-flight signal for ASC in NK cells, showing an increase in ASC area: height pulse, indicative of specking, after nigericin in vitro stimulation. **I)** Graph showing the frequency of CD56^dim^ NK cells with signs of ASC specking after nigericin stimulation. **J)** The same for CD56^bright^ NK cells stimulated with Vbp. Statistical comparisons were performed using the Wilcoxon matched-pairs signed-rank test. **K)** and **L)** Representative images of ASC expression in nigericin-stimulated monocytes CD14^+^ obtained by fluorescent imaging flow cytometry. Both patterns can be observed, diffuse ASC expression (**K**) and speck structures (**L**). **M)** Frequency of monocytes with signs of ASC specking in nigericin-stimulated cells with or without previous treatment with the dual NLRP1/NLRP3 inhibitor ADS032. The same for NK cells (CD56^+^) in **N)**, **O)** and **P)**.

Overall, our results indicate that activated gasdermin-D is inducible and functional in NK cells, particularly by the NLRP3 and NLRP1 activators nigericin and Vbp. CD56^dim^ and CD56^bright^ NK cells behave differently under pro-inflammatory stimulation. Of note, based on our mechanistic exploration assays, we believe that the non-canonical pathway leads to the activation of NLRP1/NLRP3 inflammasomes in NK cells. Alternatively, pyroptosis of NK cells activating these inflammasomes may be hindering/underestimating the detection of the canonical pathway.

### NK cells display inflammasome activation in patients with kidney disease and undergoing organ transplantation

To elucidate if NK cells participate in the orchestration and maintenance of inflammatory processes through inflammasome signaling *in vivo*, we studied this in the context of kidney disease and organ transplantation. Recent data support the hypothesis that imbalanced inflammation might be an important driver of kidney disease that may persist even after organ transplantation [23–27]. We collected PBMCs from patients with chronic kidney disease (**Supplementary Table 1**). Western blotting studies showed high protein levels of NLRP1, active caspase-1, and gasdermin-D in the blood of these patients before and during the first week after kidney transplantation (**Figure 4A and Supplementary Figure 4**), confirming the activation of inflammasomes and pro-inflammatory cell death by pyroptosis of circulating immune cells. We then specifically studied NK cells from these PBMC. Importantly, flow cytometry analyses (as represented in **Supplementary Figures 5A and B**) confirmed that nearly 100 % of NK cells in these patients expressed NLRP1 (**Figure 4B**), about 50 % expressed ASC (**Figure 4C**), and a smaller fraction NLRP3, which increased three days after transplantation (**Figure 4D**). We also detected a fraction of NK cells with signs of speck, which was enriched three days post-transplantation (**Figure 4E and Supplementary Figure 5C**). Speck levels in NK cells were much lower than those detected in monocytes, as expected (**Supplementary Figure 5D**). Additional analyses distinguishing between specking versus non-specking cells showed higher levels of NLRP1 and NLRP3 in speck^+^ cells (**Figures 4F, G, H and I**), suggesting the activation of both inflammasomes in NK cells from patients with renal dysfunction and shortly after undergoing organ transplantation.

**Figure 4.**
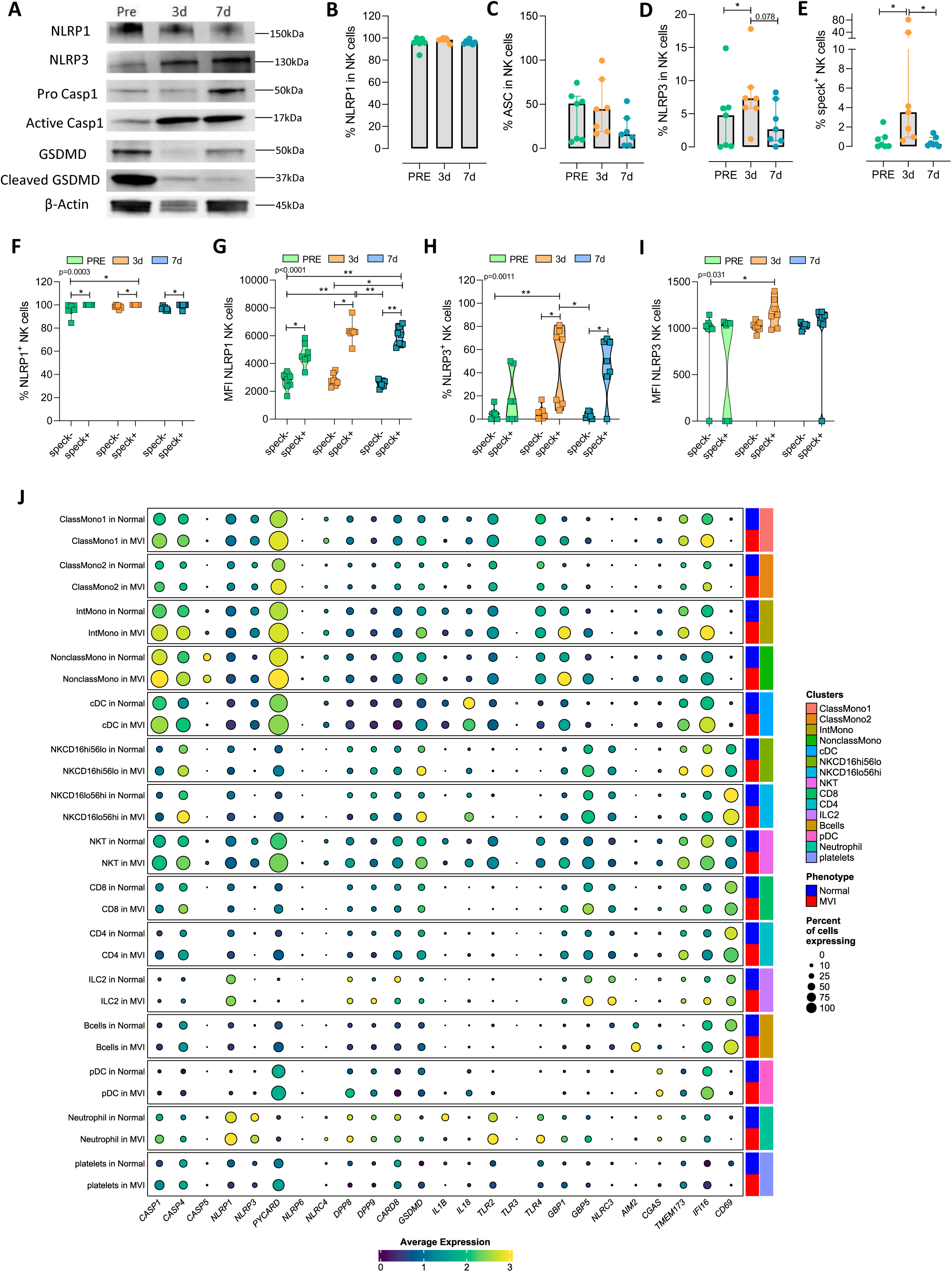
Inflammasome activation in NK cells from patients with chronic kidney disease and organ transplantation. Cells from peripheral blood samples of patients with renal disease collected at different time points before and after kidney transplantation were studied. Activation of the inflammasomes in these patients was investigated using western blotting, flow cytometry, and single-cell RNA sequencing. **A)** Western blot representative results in total peripheral blood mononuclear cells (PBMCs) from one renal-injured patient (#FJP in **Supplementary Table 1**), showing the expression of NLRP1, NLRP3, pro-active caspase-1, the active fragment of caspase-1, full-length gasdermin D, active gasdermin D, and the control β-actin, pre-transplantation and 3 and 7 days after being transplanted. **B)** Frequency of expression of NLRP1 in total CD56^+^ NK cells (n=7). **B)** Frequency of ASC. **D)** Frequency of NLRP3. **E)** Percentage of ASC specking in NK cells. Graphs represent median with range. Statistical comparisons were performed using the Wilcoxon matched-pairs signed-rank test. Graphs in **F)** and **G)** show the percentage and MFI of NLRP1 within speck^-^ and speck^+^ NK cells at different time points. The same for NLRP3 in **H)** and **I).** Graphs represent median with range. P values shown in the graphs represent ANOVA Friedman test, and asterisks denote the multiple comparisons Dunn’s test. *p<0.05; **p<0.01. Moreover, comparisons between the speck^-^ and speck^+^ NK cells at the same time points were performed using the Wilcoxon matched-pairs signed-rank test. **J)** Single-cell RNA-sequencing (scRNAseq) analysis on 12 peripheral blood samples, n=6 samples from kidney transplant recipients with a diagnosis of microvascular inflammation (MVI), and n=6 patients without MVI. We show the expression of genes implicated in the inflammasome pathways across the peripheral blood cells. cDC, conventional dendritic cell; ILC2, innate lymphoid cell; NK, natural killer cell; NKT, natural killer T cell; PBMC, peripheral blood mononuclear cell; pDC, plasmacytoid dendritic cell.

Microvascular inflammation (MVI) is a hallmark of immune-mediated injury of renal allografts, characterized by immune cell infiltration into glomerular and peritubular capillaries. Traditionally, MVI has been associated with antibody-mediated rejection (ABMR), but many MVI cases lack detectable donor-specific antibodies (DSA), suggesting the involvement of other immune mechanisms or alternative triggers such as ischemia/reperfusion and viral infections [28]. Recent studies have emphasized the role of NK cells in this lesion [29–31]. To further dissect the molecular expression patterns associated with the MVI observed in a fraction of kidney-transplanted individuals, we conducted single-cell RNA-seq analyses of PBMC samples from a previous study on kidney transplant recipients [32]. Clinical data of the patients are shown in **Supplementary Table 2**. Several circulating immune cell clusters were analyzed (**Supplementary Figure 6A and B**). A pro-inflammatory phenotype of NK cells in these individuals was confirmed, with more frequent expression of *CASPASE-1*, *CASPASE-4*, *PYCARD/ASC*, *GASDERMIN-D*, *IL-18*, *GBP1*, *GBP5*, *STING*, and *IFI16*, in both circulating CD56^dim^ and CD56^bright^ NK cells, compared to individuals lacking a microvascular inflammatory signature in the biopsy (**Figure 4J**). This was detected despite a long follow-up period after the transplant of up to 5.5 years for some of the patients participating in the study. Notably, a significant increase in the intensity of expression of caspase-4 and gasdermin-D was noticed in NK cells from patients with MVI, standing out over other immune cell subsets, and confirming non-canonical inflammasome activation and pyroptosis in these cells, even after long periods following transplantation (**Figure 4J**). Indeed, further classification of the data based on the time after transplant in *mid-* (363-730 days) and *long-term* (+731 days) showed a peak of expression of caspase-4 in NK cells at mid-term, approximately two years after receiving the organ (**Supplementary Figure 7**).

Last, we investigated the transcriptional changes occurring in NK cells at the single-cell level in kidney biopsies from transplanted patients with or without MVI, and the relationship with this outcome. Single-cell RNA-seq publicly available datasets [30] of biopsies showing MVI or not were selected as described in the Materials and Methods section. Structural compartments and several immune cell clusters of the human transplanted kidney were analyzed (**Figure 5A and B**). scRNASeq is not efficient to capture *NCAM1* expression encoding CD56, so we also used CD16 (*FCGR3A*) to distingüish CD56^dim^ and and CD56^bright^ NK subpopulations. In addition, CD16 (*FCGR3A*+) NK cells represent a subpopulation with potential to mediate antibody-dependent cell cytotoxicity (ADCC) responses, highly relevant in the context of ABMR [31]. Clinical data from patients is summarized in **Supplementary Table 3**. Similar to the results in peripheral blood, higher levels of caspase-4 were found in CD16 (*FCGR3A*+)-expressing NK cells (**Figure 5C-D**), as well as higher NLRP1 and lower levels of DPP8 and DPP9 in both CD16^+^ and CD16^-^ NK cells in biopsies with inflammatory lesions (**Figure 5C**). We did not observe an increase in gasdermin-D in renal NK cells, which could be attributed to the cells undergoing pyroptosis more rapidly and thus dying. This inflammatory flux might promote the migration of lymphocytes from circulation to inflamed kidneys and explain the higher frequencies of immune effectors such as NK cells and T cells in biopsies showing MVI (**Figure 5E**).

**Figure 5.**
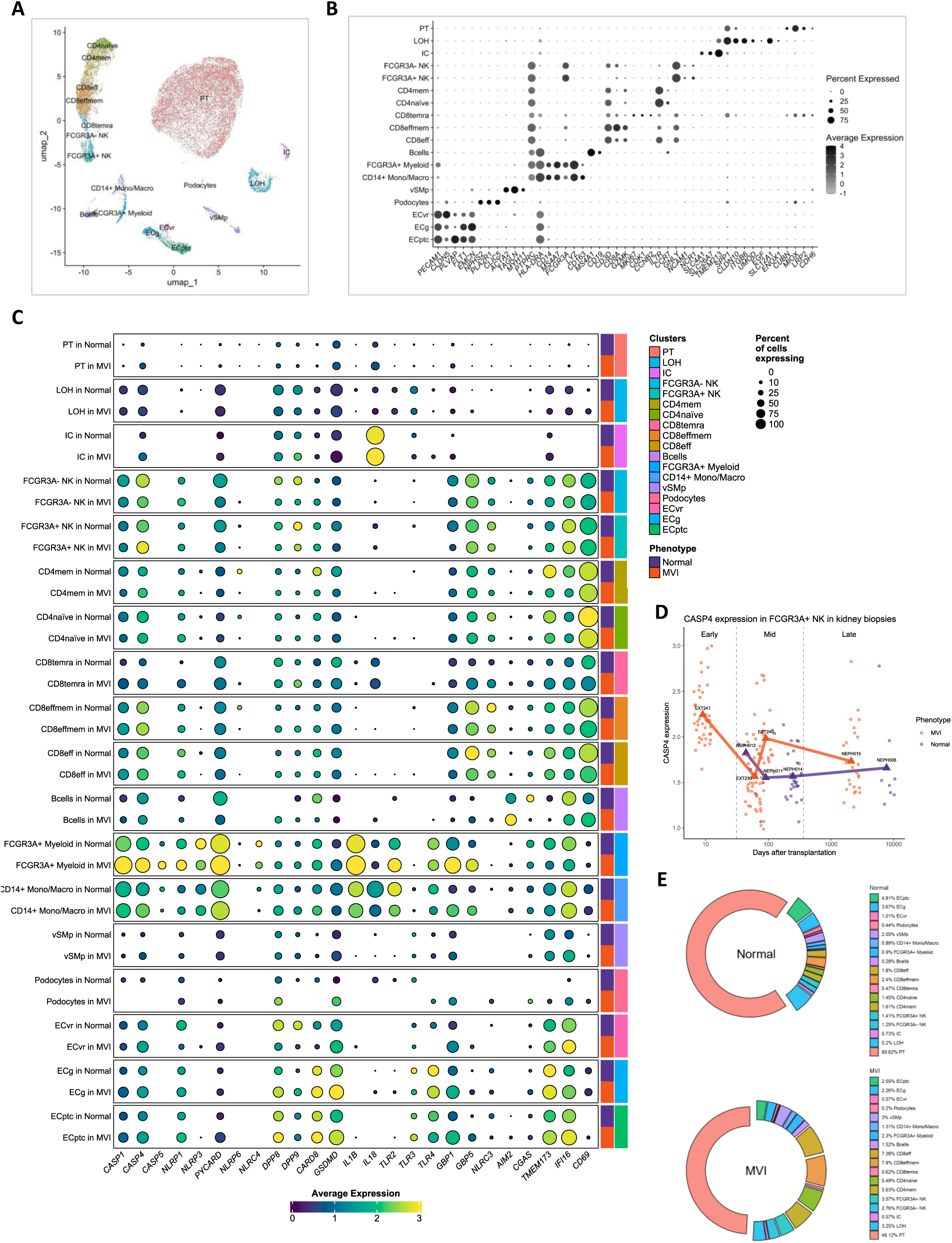
**A)** Analysis of scRNAseq data from kidney allograft biopsies (*n* =9 biopsies from 4 transplant recipients with a diagnosis of microvascular inflammation (MVI), and 5 patients without MVI (normal). Uniform Manifold Approximation and Projection (UMAP) plot of 20,607 cells passing QC filtering. The main kidney cell types are represented, including loop of Henle (LOH), podocytes, vascular smooth muscle and pericytes (vSMp), proximal tubule (PT), intercalated cells (IC), three endothelial cell subsets comprising vasa recta (ECvr), glomerular (ECg) and peritubular capillaries (ECptc), myeloid cells, and lymphoid cells such as FCGR3+ and FCGR3-NK cells. **B)** Dot plot showing average gene expression values of canonical lineage markers (log scale) and percentage of major cell types represented in the dataset and UMAP plot. **C)** Graph showing the expression of genes implicated in the inflammasome pathways across the different cell types analyzed and comparing between “normal” biopsies and those showing microvascular inflammation (MVI). **D)** Caspase-4 expression in *FCGR3*+ NK cells according to time after transplantation in kidney biopsies (Early: 0-31 days; Mid: 32-365 days; Late: >366 days). **E)** Relative proportions of the different cell types identified in the kidney transplant biopsies.

## Discussion

In this study, we shed light on the biology of inflammasomes in human NK cells, as dedicated studies are lacking. These multiprotein complexes are involved in innate defense and inflammatory responses to pathogens. Similarly, NK cells constitute a first line of defense against aberrant targets. However, whether inflammasomes encompass one of their multiple immune mechanisms of action and how it may impact health are unknown. Inflammasomes are composed of sensors, adaptors, and effectors components, and their activation triggers the release of inflammatory cytokines and pyroptosis [33]. Here, we show the expression of transcripts encoding several inflammasome-forming proteins in NK cells distributed across different tissues. Sensors of several inflammasomes, including the members of the NALP family NLRP3 and NLRP1 were found expressed. Consistent with a prior report [34], we found a high constitutive expression of NLRP3. We also noticed high levels of NLRP1 and the adaptor protein ASC in NK cells. More importantly, we prove here that inflammasomes are functional in these cells and non-canonical activation can occur.

Several activators of the NLRP3 and NLRP1 inflammasomes have been identified, being reported potassium ion (K^+^) efflux as one of the most potent in the case of NLRP3 [35], and recently also proven to activate NLRP1 [36], and inhibition of the serin dipeptidyl peptidases DPP8 and DPP9 for NLRP1 activation [20, 21]. Here, we used Nigericin, a toxin derived from *Streptomyces hygroscopicus* that facilitates H^+^/K^+^ anti-port across cell membranes and thereby causes potassium efflux [37], and Vbp, an inhibitor of the inhibitory interaction between DPP8/DPP9 and NLRP1 [38], to assess the functionality of both inflammasomes in NK cells. Our results revealed changes that are compatible with inflammasome activation, including prominent signs of pyroptosis. Moreover, our experiments seemed to indicate that the components of the NLRP3 and NLRP1 inflammasomes are more likely to undergo a reorganization rather than upregulation upon stimulation with nigericin or Vbp. Besides, we found differences in the response of CD56^dim^ and CD56^bright^ NK cells to both compounds. This is not completely unexpected given the high plasticity and diversity in phenotypes and functions of NK cells [2, 3]. Regarding the pore-forming protein gasdermin D, capable of inducing pyroptosis, its active form was detected in CD56^dim^ NK cells treated with nigericin and Vbp. LDH levels were also increased in the culture of these cells, particularly in those treated with nigericin. Of note, although we could not detect active gasdermin D in CD56^bright^ cells, a notable release of LDH by these cells was observed. The release of LDH is considered a surrogate of cell pyroptosis as large proteins are usually filtered out in the absence of cell death [39]. Thus, it may be plausible that CD56^bright^ NK cells undergo more aggressive pyroptosis upon *in vitro* activation of the NLRP3/NLRP1 inflammasomes, hindering the detection of active gasdermin-D. Also, the low numbers of these cells in the blood, which usually represent less than 10% of circulating NK cells [40], may pose a technical challenge for the detection of proteins. Alternatively, other forms of cell death could be involved. Different frequencies and proportions of NK cell subpopulations can be found in tissues like the bone marrow, spleen, lung, liver, lymph nodes, tonsils, and intestines [3]. Thus, further studies with different tissue samples, NK subpopulations, and inflammasome-triggering agents or conditions could add valuable insight into the biology of these lymphocytes and the inflammatory field.

To our knowledge, this is the first report clearly showing the inflammasome activation and pyroptosis in human NK cells. Nevertheless, prior results in mice support our results, showing that burn injury and radiation exposure induce the activation of caspase-1 in many cell subsets including NK cells [41, 42]. Importantly, Fong et al also showed that the human pathogen group B *Streptococcus* directly interacts with NK cells and suppresses a pyroptotic-like response via the interaction with siglec-7 [43]. In this regard, our results did not show a typical inflammasome response after the LPS-ATP stimulation of NK cells. However, this could be explained by the low surface expression of TLR4 in these cells [44]. Thus, we do not rule out the possibility that direct bacterial infection might activate some of the inflammasomes in NK cells. In fact, our transcriptomic analysis showed that many other inflammasome sensors are transcriptionally expressed in these cells, including NLRP6, NLRC4 or AIM-2, although if they are functional remain unknown and could be more suitable after direct infections. Moreover, this evidence supports the notion that in vivo different microenvironments and pathologic threads could trigger different inflammasome routes in NK cells. Thus, the results presented here provide new insight into the biology of NK cells that might support the role of these lymphocytes in different contexts via inflammasome activation. Elucidating when these responses take place and whether they are beneficial or harmful could help in the identification of new targets in inflammatory disorders.

Inflammasome activity has been associated with chronic kidney disease and kidney allograft rejection, particularly linked to AIM-2, NLRP3, and more recently NLRP1 [23–27]. Here, we observed signs of NLRP3 and NLRP1 inflammasome activation as well as activation of gasdermin-D in circulating immune cells, including NK cells, in patients with end-stage kidney disease before and after receiving a donor’s kidney. Previously, it has been shown that post-transplantation NK cells subsets can change even at the peripheral level, with variations in number and phenotype, and that they are enriched in biopsies from patients with antibody-mediated rejection (ABMR) and associated with inflammatory lesions in the microvascular compartment [30, 45].

Herein, we show the acquisition of a proinflammatory phenotype in NK cells shortly post-transplantation, characterized by higher levels of NLRP1, NLRP3, and ASC, which persist long-term marked by upregulated caspase-4. This suggests that different inflammatory milieus in vivo could induce the activity of inflammasomes not only in classically studied subsets such as monocytes and macrophages but also in innate lymphoid cells such as NK cells. Until now, two mechanisms have been considered the main drivers for graft rejection in transplanted patients, T-cell-mediated rejection, and/or antibody-mediated rejection. In both, NK cells have been shown to cooperate, through the maturation of dendritic cells, activation of T cells, the triggering of missing-self activation due to mismatches in HLA class I loci and NK inhibitory killer immunoglobulin-like receptors (KIR), and mediating a pathogenic ADCC response triggered by anti-HLA antibodies in CD56^dim^ CD16-expressing NK cells [46]. Our results add evidence that NK cells might also contribute to pathogenic inflammation in renal injured patients through pro-inflammatory activity via inflammasome signaling. Although larger studies and longer follow-up periods are needed, our results are tempting to speculate that inflammasome is a pathway by which NK cells might influence renal disease and graft outcomes.

Recent studies have associated the activity of NK cells, particularly by Fc-receptor mediated activation and ADCC, with graft injury patterns and ABMR, the principal process associated with MVI [29–31, 47, 48]. This scenario requires the production of DSA able to mediate ADCC. Nevertheless, many MVI cases lack detectable DSA [49, 50], suggesting the existence of multiple and alternative immune mechanisms responsable for this inflammation. Both, DSA-positive (ABMR) and DSA-negative MVI have been associated with similar NK cell burden [51], so they might be impacted by similar NK cell activation mechanisms [28]. Some mechanisms have been proposed, including the “missing-self” activation of NK cells, the IFN-γ response, and a particular receptor repertoir potentially driving strong immune responses, although they do not fully explain the diverse and complex nature of MVI [28]. Given the multifaceted role NK cells could play in MVI, targeting these cells and their effector mechanisms could help reduce MVI. Indeed, in a recent phase 2 trial conducted in late HLA DSA-positive ABMR patients, the anti-CD38 antibody “felzartamab”, showed promising results, substantially reducing MVI scores and with most patients converting to “no ABMR” or an “inactive” ABMR phenotype [52]. Interestingly, this effect was accompanied by marked decreases in the number of circulating NK cells and particular NK cell transcript sets [53]. Based on our results, it could be hypothesized that NK cells are also relevant immune sensing effectors and orchestrators of a pro-inflammatory response through inflammasome activation, potentially recruiting other cell subsets such as monocytes/macrophages, neutrophils and T cells to the site of inflammation, and thus sustaining this response. In this case, targeting inflammasomes and preventing pyroptosis could represent an effective mechanism to reduce MVI. In this sense, several inhibitors of gasdermin-D are being tested [54]. Moreover, in light of our results, both NLRP1 and NLRP3 inflammasomes could be active and associated with MVI. Unfortunately, up to date, only one dual-action molecule inhibitor targeting both the NLRP1 and NLRP3 inflammasomes has been reported but it has not been clinically approved yet [55]. Thus, the development of dual inflammasome-targeting pharmacological approaches could be highly beneficial in the context of MVI and organ rejection.

Finally, we would light to emphasize that NK cells might play a role in several other inflammatory diseases such as ischemic stroke, sepsis, or cancer, where persistent inflammasome activity and activation of gasdermin D have been observed to cause undesired effects including organelle dysfunction, cell lysis, and persistent release of cytokines [14]. Also, it will be relevant to determine other contexts where inducing inflammasome activity of NK cells could be beneficial.

In conclusion, our findings underscore the importance of including NK cells in future studies investigating inflammasome functionality in different clinical contexts.

## Acknowledgments

M.D.C is supported by PI21/01656 grant from Instituto de Salud Carlos III, Spain, EMC21_00033 EMERGIA program from Junta de Andalucia, AthenaDAO, PRF 2021-78 from Progeria Research Foundation, and grant CNS2024-154939 from the Ministerio de Ciencia, Innovación y Universidades, Agencia Estatal de Investigación. A.A.G was supported by the Spanish Secretariat of Science and Innovation post-doctoral contract Juan de la Cierva (FJC2021-047304-I). The funders had no role in study design, data collection and analysis, the decision to publish, or the preparation of the manuscript.

## Author contributions

Conceptualization: A.A.G, M.D.C; Methodology: A.A.G, I.M.Z, AV, BL; Investigation: A.A.G, JM.S.R, AT, JK, M.D.C; Resources: A.A.G, J.L.P, R.V.M, MN, AM, M.D.C; Visualization: A.A.G, I.M.Z, AV, AT, JK, BL, M.D.C; Supervision: A.A.G, M.D.C; Funding acquisition: M.D.C; Writing – Original Draft: A.A.G; Writing –Review & Editing: A.A.G, M.D.C.

## DECLARATION OF INTEREST

The authors declare no competing interest.

---

**Supplementary Table 1**. Clinical data of patients with renal dysfunction included in the study, and sex and age of the organ donors.

**Supplementary Table 2**. Clinical data of patients included in the single cell transcriptomic analysis of peripheral blood samples from kidney transplant recipients, comparing clinical parameters between those with signs of MVI and those lacking MVI signature.

**Supplementary Table 3**. Clinical data of patients included in the scRNAseq analysis from kidney allograft biopsies.

## MATERIALS AND METHODS

### Human Samples

In this study, we used primary cells obtained from blood samples from healthy donors or patients with kidney disease and undergoing transplantation. Peripheral blood mononuclear cells (PBMCs) were obtained from the Hospital Puerta del Mar in Cádiz, Spain. Study protocols were approved by the corresponding Ethical Committees of the Hospital. All subjects recruited for this study were adults who provided written informed consent. Information on renal disease parameters from affected patients is summarized in Supplementary Table 1. Gender and age are also indicated in Table S1 unless not available. Since we utilized samples from donors of different genders in our experiments, we can conclude that the results reported here apply to both men and women. PBMCs were obtained by Ficoll-Paque density gradient centrifugation. PBMCs were cultured in RPMI medium (Gibco) supplemented with 10% Fetal Bovine Serum (Gibco), 1% streptomycin-penicillin (Thermo Fisher, Waltham, MA, USA, 11548876) (R10 medium), and maintained at 37°C in a 5% CO_2_ incubator, when needed. NK cells were isolated from PBMCs by FACS (fluorescence-activated cell sorting) (MELODY sorter) or using a commercial kit (MagniSort™ Human NK cell Enrichment; eBioscience). The purity of the cells was always higher than 90%.

### Transcriptomic analysis of a publicly available RNAseq data

We analyzed publicly available RNA-seq datasets. Data are available from Gene Expression Omnibus (GEO) corresponding to the study with accession number GSE133383. This study provides valuable transcriptomic data of two subpopulations of NK cells, CD56^dim^ and CD56^bright^, in tissues difficult to obtain [3]. R version 4.1.2 was used to perform all bioinformatic analyses. Count values were imported and processed using edgeR [https://academic.oup.com/bioinformatics/article/26/1/139/182458?login=false]. Expression values were normalized using the trimmed mean of M values (TMM) method and lowly-expressed genes (<1 counts per million) were filtered out. Differentially expressed genes were identified using linear models (Limma-Voom) [https://academic.oup.com/nar/article/43/7/e47/2414268], and P values adjusted for multiple comparisons by applying the Benjamini-Hochberg correction. Heatmaps were generated using the heatmap3 package [https://github.com/slzhao/heatmap3]. Glimma [https://github.com/Shians/Glimma] was used for interactive visualizations. Functional gene annotation was performed using DAVID [https://david.ncifcrf.gov].

### Single-cell transcriptomic data analysis

Publicly available datasets corresponding to kidney allograft biopsies showing MVI or not were selected from E-MTAB-12051 [30] as followed: EXT217 = Normal, EXT230 = MVI, EXT240 = MVI, EXT241 = MVI, NEPH006 = Normal, NEPH011 = Normal, NEPH012 = Normal, NEPH014 = Normal, NEPH015 = MVI. Publicly available datasets corresponding to PBMCs from kidney transplant recipients showing MVI or not at the time of blood sample (Supplementary Table 2) were gathered from E-MTAB-11450 [32]. Analysis was performed as previously described [30, 32], notably for UMAP, grey scale dot plots and proportions. Viridis colored clustered dot plots were generated using the DotPlot_Heatmap function from the R package RightOmicsTools: Varin, A.: An R Package Providing Complementary Tools for the Manipulation and Visualization of Single Cell RNA-Seq Data. Preprint at https://doi.org/10.5281/zenodo.12518909 (2025).

### Human NK cell isolation by FACS

For Fluorescence Activated Cell Sorting (FACS), 80 million PBMCs were stained with LIVE/DEAD AQUA viability (Invitrogen) for 20 minutes at RT. After washing with staining buffer (PBS 3% FBS), cells were surface stained with anti-CD56-FITC (B159, Becton Dickinson), anti-CD3-PE-Cy7 (SK7, Becton Dickinson), anti-CD16-BV786 (3G8, Becton Dickinson) and anti-CD4-APC (RPA-T4, Becton Dickinson) antibodies for 20 minutes at RT. Cells were then washed with staining buffer and immediately sorted using the MELODY Cell Sorter. We sorted the live populations CD4^-^CD3^-^CD56^dim^ (CD56^dim^ NK cells) and CD4^-^CD3^-^CD56^bright^ (CD56^bright^ NK cells). The purity of the cells was >90% in all cases.

### In vitro inflammasome activation assays

For in vitro pro-inflammatory stimulation, NK cells were treated with 1µg/ml LPS (Sigma-Aldrich, St. Louis, MI, USA, L4391-1MG) for 4 hours and 5 mM ATP (Santa Cruz Biotechnology, Santa Cruz, CA, USA, sc-214507A) for 30 mins. NK cells were also stimulated with 1 µM Val-boroPro (Sigma-Aldrich, St. Louis, MI, USA, 5314650001) for 4 hours or 10 µM nigericin for 45 mins. As the number of isolated NK cells varied between donors, we maintained the conditions of the cell culture, stimulating the cells at a concentration of approximately 0,1M/ml, in 24-well plates for the CD56^dim^ subpopulation and 96-well plates for the CD56^bright^ subpopulation. After the incubation, we collected culture supernatants for LDH and cytokine measures, and cells for proteomics analyses by flow cytometry or western blot. Controls consisting of cells previously treated with the dual NLRP1/NLRP3 inflammasome inhibitor ADS032 (100 µM, MedChemExpress) were included.

### Flow cytometry

For studying the protein expression, before and after pro-inflammatory stimulation, NK cells were stained with LIVE/DEAD AQUA viability (Invitrogen) for 20 minutes at room temperature (RT). After washing once with staining buffer (1X PBS 3% FBS), cells were surface stained with anti-CD56-FITC (B159, Becton Dickinson), and anti-CD3-PE-Cy7 (SK7, Becton Dickinson) antibodies for 20 mins at RT. Next, we performed a washing step, and cells were fixed and permeabilized with Fixation/Permeabilization Solution (Becton Dickinson) for 20 minutes at 4°C and then washed with BD Perm/Wash buffer. After, cells were stained with rabbit anti-NLRP1 (A16212 ABclonal) for 20 mins at RT, washed, and detected by staining with an anti-rabbit secondary AF750 (ab175735, Abcam) antibody for an additional 20 mins at RT. Another washing step was performed and staining with anti-NLRP3-AF700 (768319, R&D Systems) and anti-ASC-PE (HASC-71, Biolegend) was carried out for 20 minutes at RT. Finally, cells were washed with BD Perm/Wash.

We also studied the expression of the inflammasome components in NK cells and monocytes from kidney-injured patients, before and after being subjected to kidney transplantation. PBMCs from these patients were stained with LIVE/DEAD AQUA viability (Invitrogen), and then, with anti-CD56-FITC (B159, Becton Dickinson), anti-CD3-PE-Cy7 (SK7, Becton Dickinson), anti-CD4-BV605 (RPA-T4, Becton Dickinson), anti-HLA-DR-PE-Da594 (L243, Biolegend), and anti-CD14-APC-H7 (M5E2, Becton Dickinson) antibodies. Cells were subsequently fixed and permeabilized with Fixation/Permeabilization Solution (Becton Dickinson) and intracellularly stained with anti-NLRP1-AF647 (vwr), anti-NLRP3-AF700 (768319, R&D Systems) and anti-ASC-PE (HASC-71, Biolegend). Samples were acquired on a CELESTA flow cytometer, and data was analyzed using FlowJo V10 software. Gating was performed according to the different FMO controls.

### Western blot assays

Western blotting was performed using standard methods. Total PMBCs or NK cell extracted proteins were used for standard protein electrophoresis and Western blot assays. Gel electrophoresis was performed using 4–20% Mini-PROTEAN® TGX Stain-Free™ Protein Gels, Biorad at 200V in Tris-Glycine-SDS buffer for 40 mins. Protein transfer was made using a TurboTransfer (Biorad, Hercules, CA, USA) at 25V for 7 mins. After transferring the proteins to 0.45 µM nitrocellulose membranes (Biorad, Hercules, CA, USA), these were incubated for 1 hour in BSA 5% in PBS-Tween20 0.05% and then overnight at 4°C with primary antibodies at 1:1,000 dilution. Then washed twice with PBS-Tween20 and incubated with the corresponding secondary antibody coupled to horseradish peroxidase diluted 1:10,000 for 1h at RT. Protein loading was checked using stain-free gel activation and tubulin protein amount. Stripping was not used.

The following primary antibodies were used: NLRP1 (ABclonal, A16212), NLRP3 (ABclonal, A5652), Caspase-1 (Cell signaling, 3866S), Gasdermin D (Santacruz, sc-393581), Caspase-4 (F4T9L) (Cell signaling, 42264T), GADPH (Cell signaling, 5174S). Anti-rabbit or anti-mouse IgG secondary antibody from Calbiochem was used.

### Analysis of cytokine and immune factor secretion

Concentrations of the cytokines and molecules GM-CSF, IFN-γ, IL-1β, IL-6, IL-10, TNF-α, IL-18, IFN-α, and ICAM-1 were quantified in 50 μL of supernatant from stimulated NK cells using a bead-based multiplex immunoassay (ProcartaPlex; Invitrogen) according to the manufacturer’s recommendations. Measurements were performed using a Luminex Intelliflex instrument (ThermoFisher Scientific) and analyzed using a standard curve for each cytokine.

### LDH cytotoxicity assay

Cell death was measured by lactate dehydrogenase (LDH) release in the supernatant following the manufacturer’s instructions (abcam). In these experiments, cells were plated and stimulated in RPMI 1640 without FBS to not interfere with the assay.

### Fluorescent flow cytometry imaging for ASC specking analyses

Isolated NK cells and monocytes stimulated with Nigericin (10 µM, 50 minutes) with and without previous treatment with the NLRP1/NLRP3 dual inhibitor ADS032 (100 µM, 45 mins before adding Nigericin) were stained in 100 µl of staining buffer (PBS 3% FBS) with CD14-PE-Cy7 (HCD14, Biolegend), CD3-AF488 (SP34-2, BD), CD56-BV421 (NCAM16.2, BD). Cells were then washed and fixed and permeabilized with Fixation/Permeabilization Solution (Becton Dickinson) for 20 minutes at 4°C. After a wash with BD Perm/Wash buffer, cells were intracellularly stained with ASC-PE (HASC-71, Biolegend), and NLRP1-AF647 (vwr) or NLRP3-AF700 (768319, R&D Systems) antibodies. Cells were acquired with an AMNIS ImageStreamX imaging flow cytometer (Merck), and data was analyzed using IDEAS v6.1 software.

### Quantification and statistical analysis

Statistical analyses were performed with Prism software, version 6.0 (GraphPad). A P value <0.05 was considered significant. The statistical details for the different experiments can be found in each figure legend.

---

**Supplementary Figure 1.**
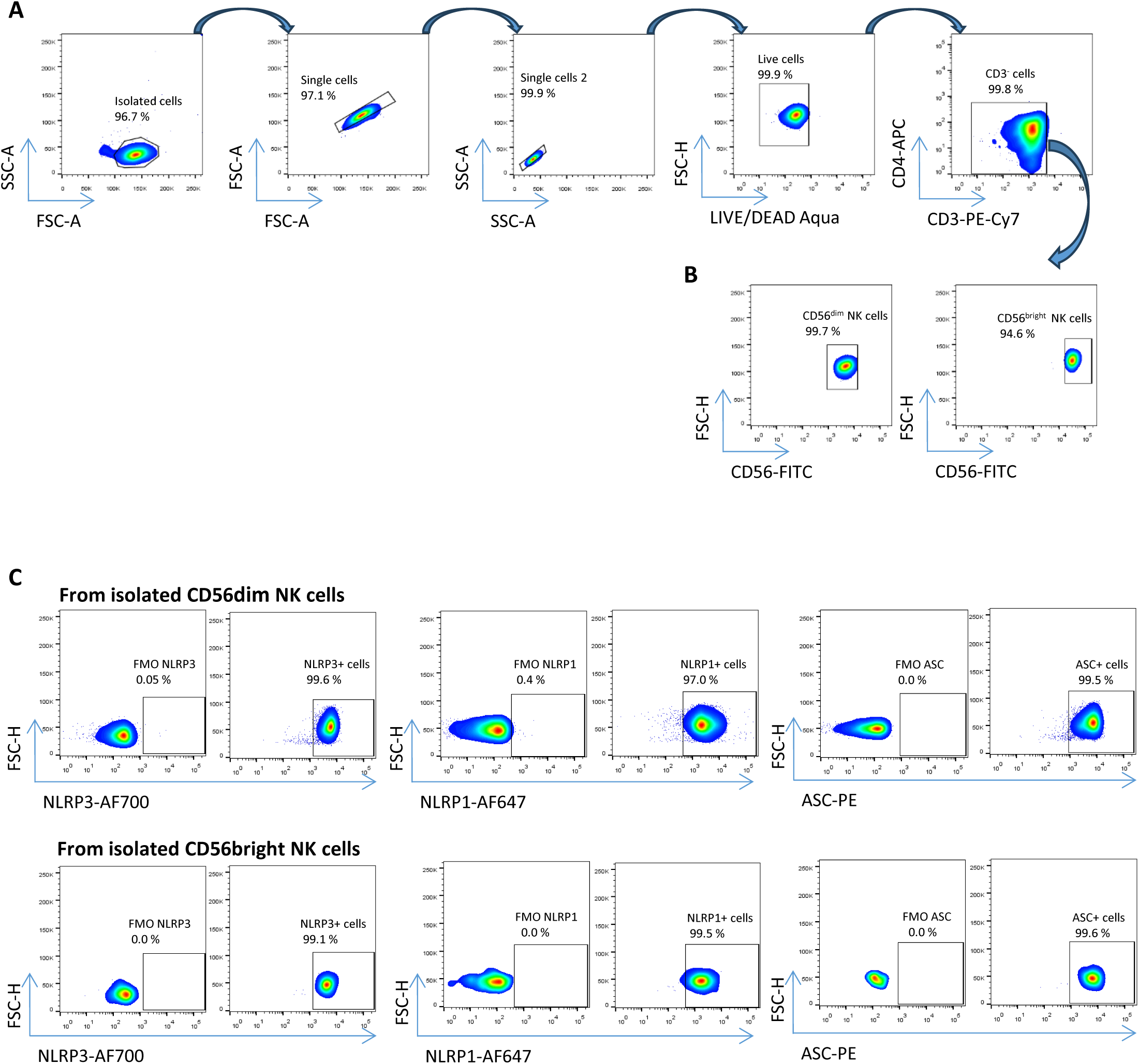
**A)** Example of the gating strategy for the identification of NK cells. **B)** Example of the purity of the CD56^dim^ and CD56^bright^ NK cells after cell sorting. **C)** Flow-cytometry plots showing a representative example for the expression of NLRP1, NLRP3, and ASC in CD56^dim^ and CD56^bright^ NK cells, and the corresponding Fluorescence Minus One (FMO) control for the proper gating.

**Supplementary Figure 2.**
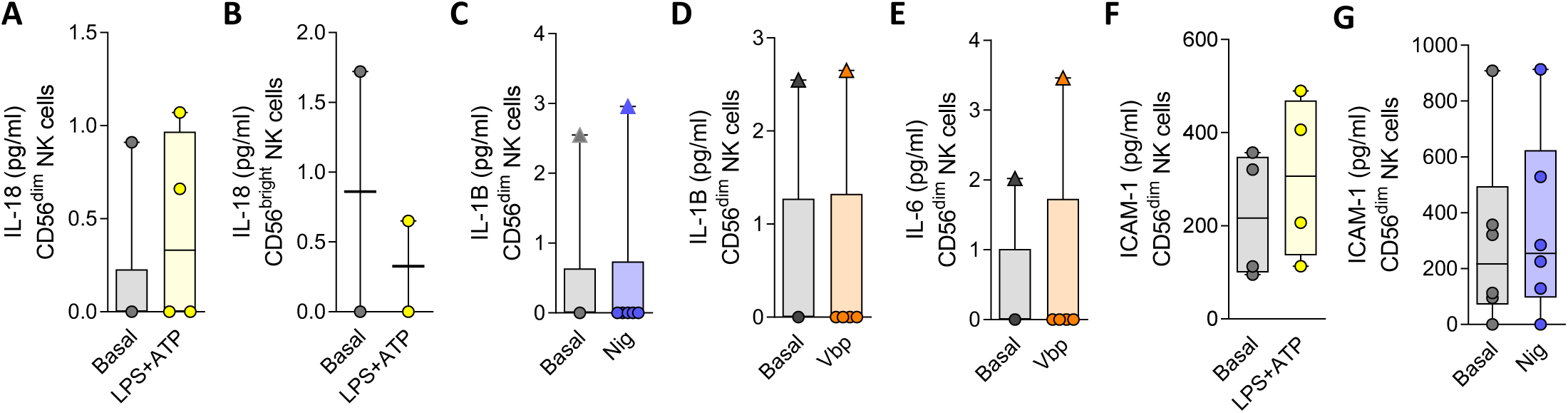
**A)** Concentration of IL-18 after the stimulation of CD56^dim^ NK cells with LPS-ATP. **B)** The same for CD56^bright^ NK cells. **C)** Concentration of IL-1β after stimulation of CD56^dim^ NK cells with nigericin, or **D)** with Vbp. **E)** Concentration of IL-6 in cell cultures of CD56^dim^ NK cells stimulated with Vbp, and **F)** concentration of ICAM-1 after LPS-ATP or **G)** nigericin stimulus.

**Supplementary Figure 3.**
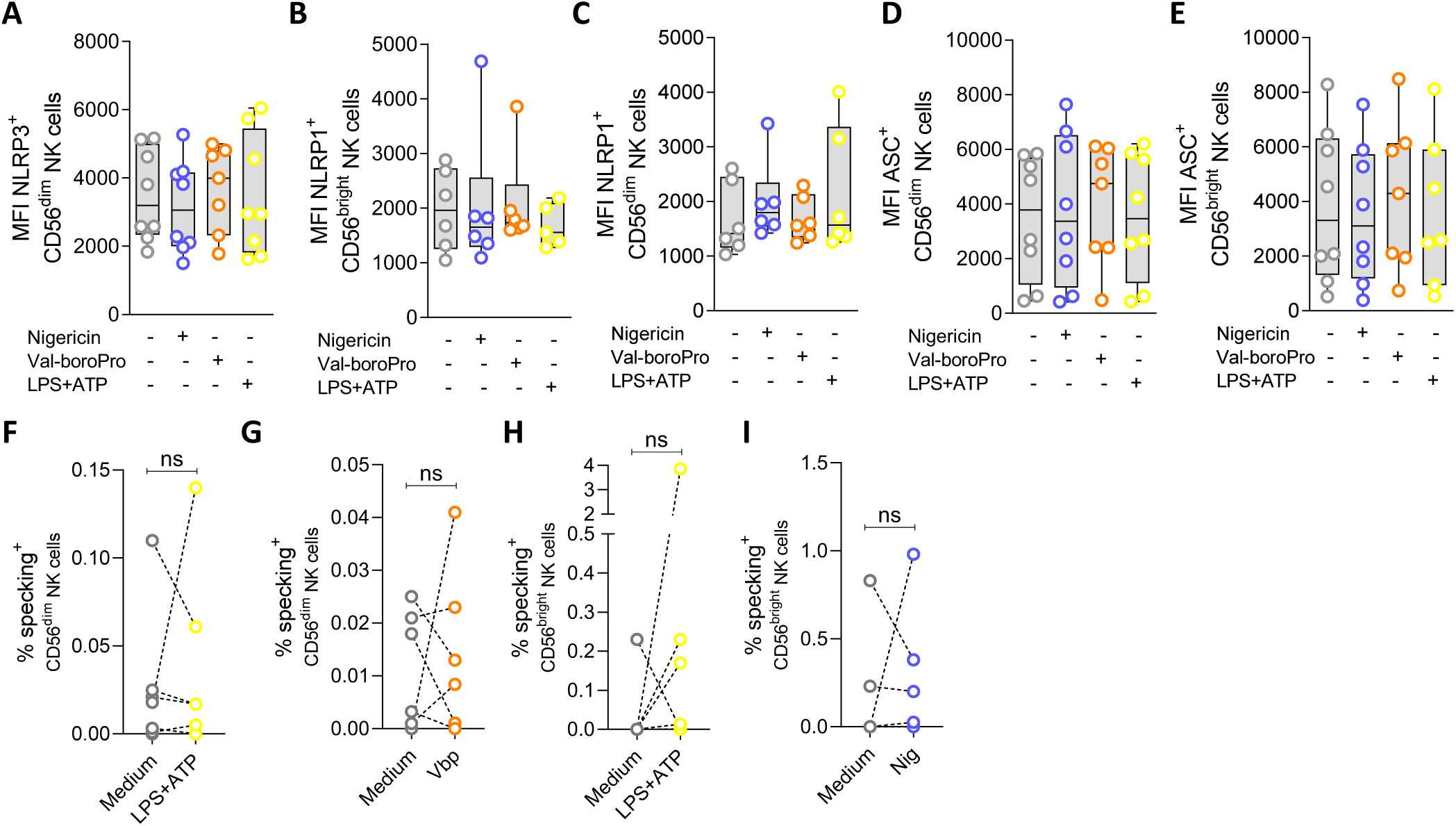
**A)** to **E)** Mean Fluorescence Intensity (MFI) signal for NLRP1, NLRP3 and ASC in separate cultures of CD56^dim^ and CD56^bright^ NK cells basally and after pro-inflammatory stimulation with LPS-ATP, nigericin or Vbp. Graphs represent median with range. **F)** to **I)** Frequency (%) of specking cells within CD56^dim^ or CD56^bright^ NK cells after stimulation with LPS-ATP, nigericin or Vbp. Statistical comparisons were performed using the Wilcoxon matched-pairs signed-rank test.

**Supplementary Figure 4.**
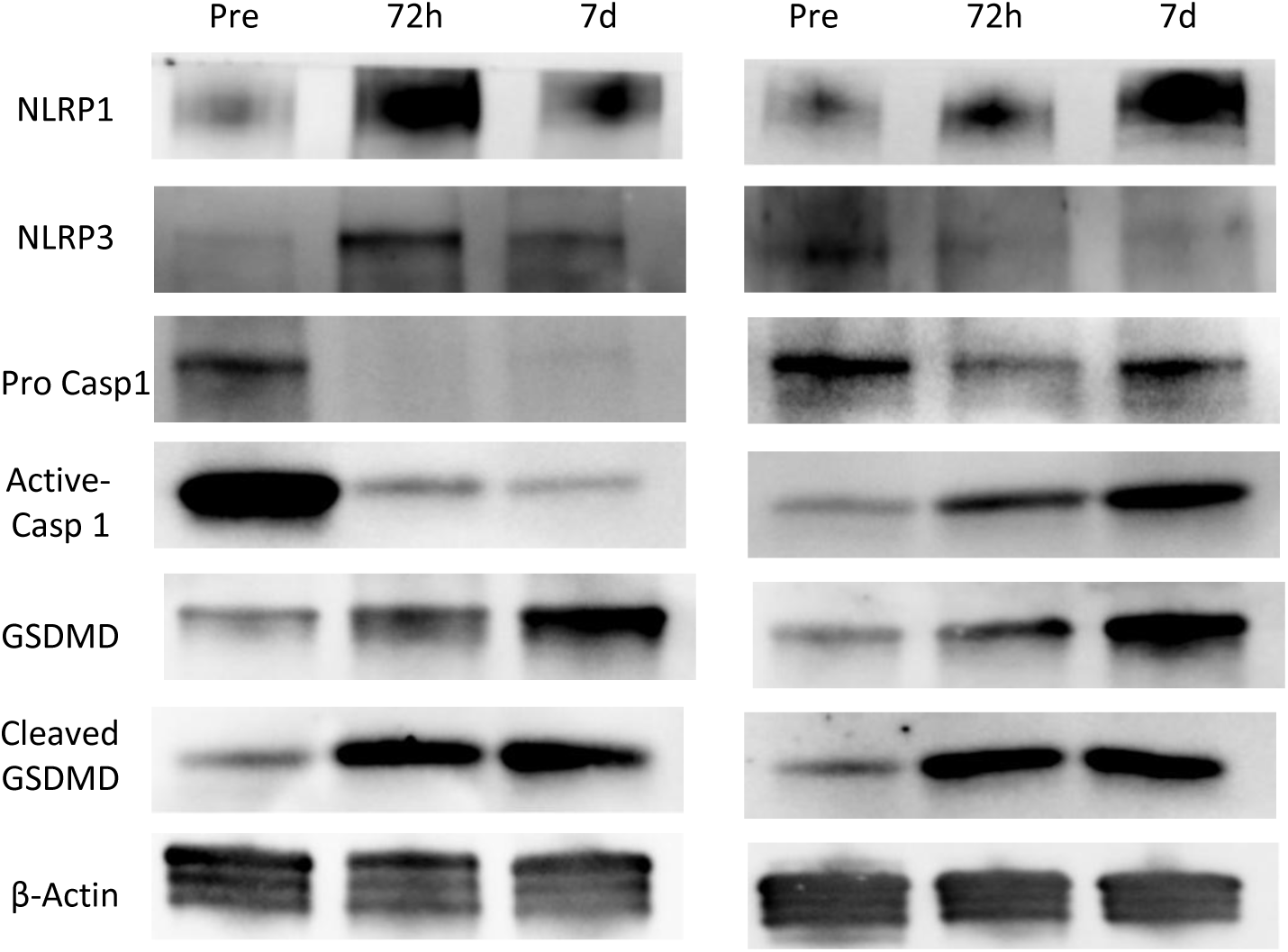
Results from the analysis by western blot of the expression of several inflammasome proteins in PBMCs from two patients with renal dysfunction (#MCB and #MFS in **Supplementary Table 1**). This figure shows the expression of NLRP1, NLRP3, pro-active caspase-1, the active fragment of caspase-1, full-length gasdermin D, active gasdermin D, and the control β-actin, pre-transplantation and 3- and 7-days post-transplant.

**Supplementary Figure 5.**
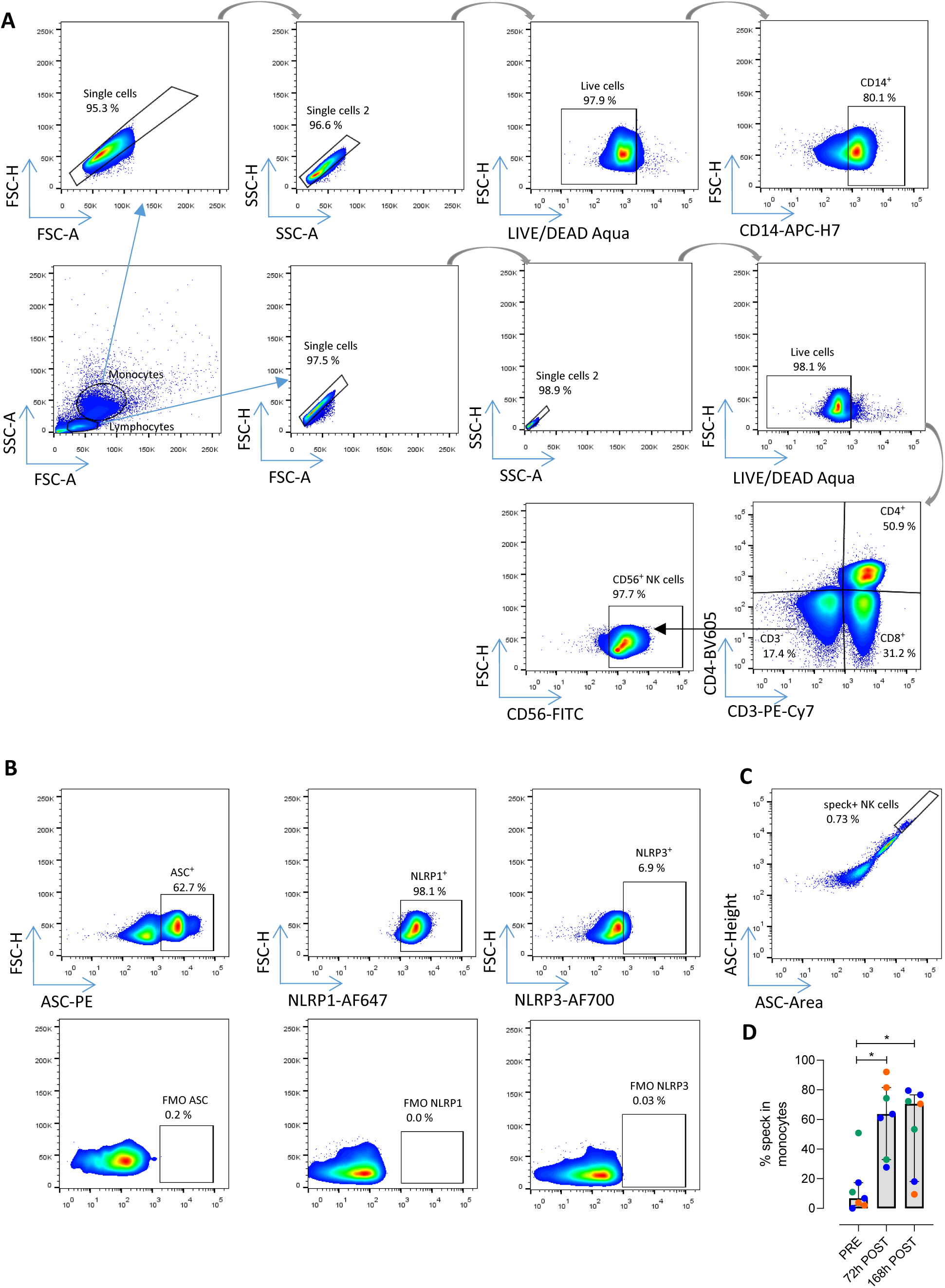
**A)** Representative example of the gating strategy used for the identification of monocytes and NK cells in samples from patients with kidney dysfunction. **B)** Representative plots of the expression of ASC, NLRP1 and NLRP3 in NK cells from these patients. FMO controls are shown. **C)** Representative plot of the specking levels within NK cells from renal-injured patients by the time-of-flight signal for ASC, showing an increase in ASC area: height pulse. **B)** Frequency of speck^+^ monocytes in the samples from patients with renal dysfunction pre- and post-transplant (n=7). The graph represents median with range and statistical comparisons were performed using the Wilcoxon matched-pairs signed-rank test.

**Supplementary Figure 6.**
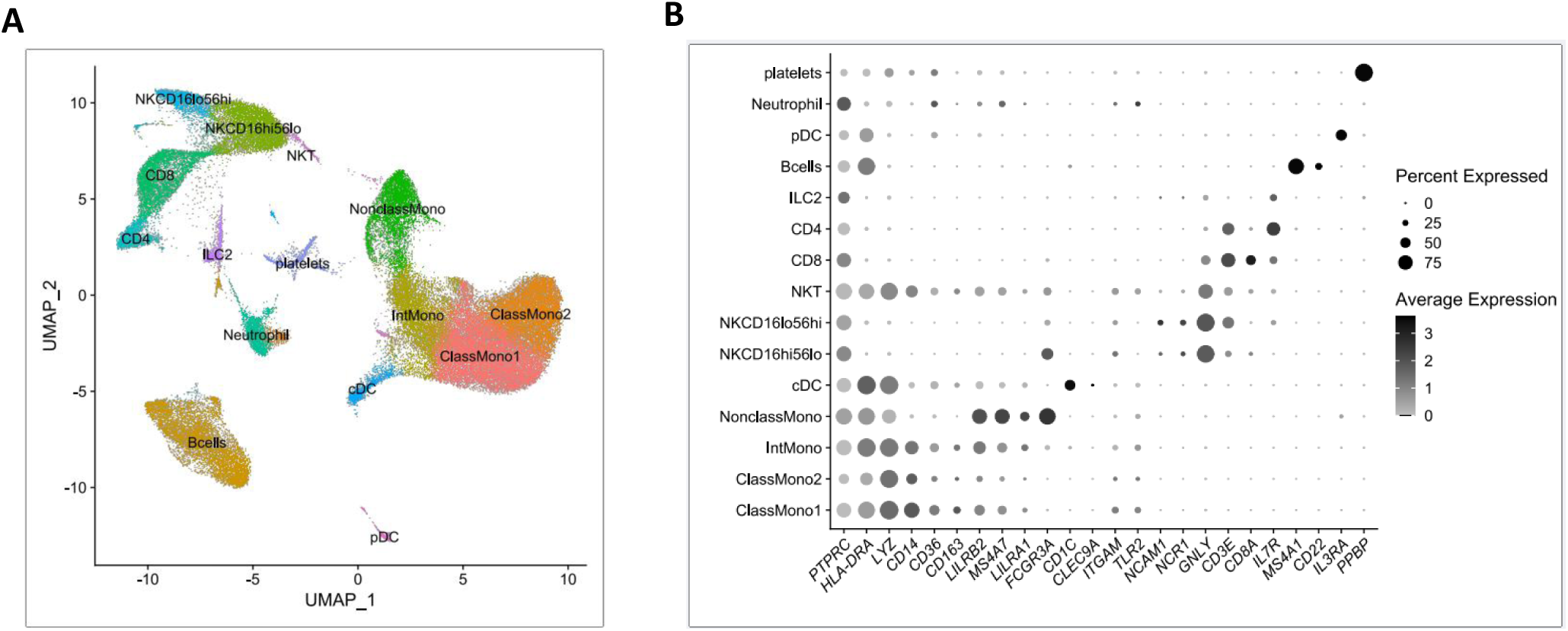
Single-cell RNA-sequencing (scRNAseq) analysis on 12 peripheral blood samples, n=6 samples from kidney transplant recipients with MVI, and n=6 patients without MVI. **A)** Uniform Manifold Approximation and Projection (UMAP) representing several circulating immune cell types, including lymphoid and myeloid cells. **B)** Dot plot showing average gene expression values of canonical lineage markers (log scale) and percentage of major cell types represented in the dataset (Figure 4J) and UMAP plot.

**Supplementary Figure 7.**
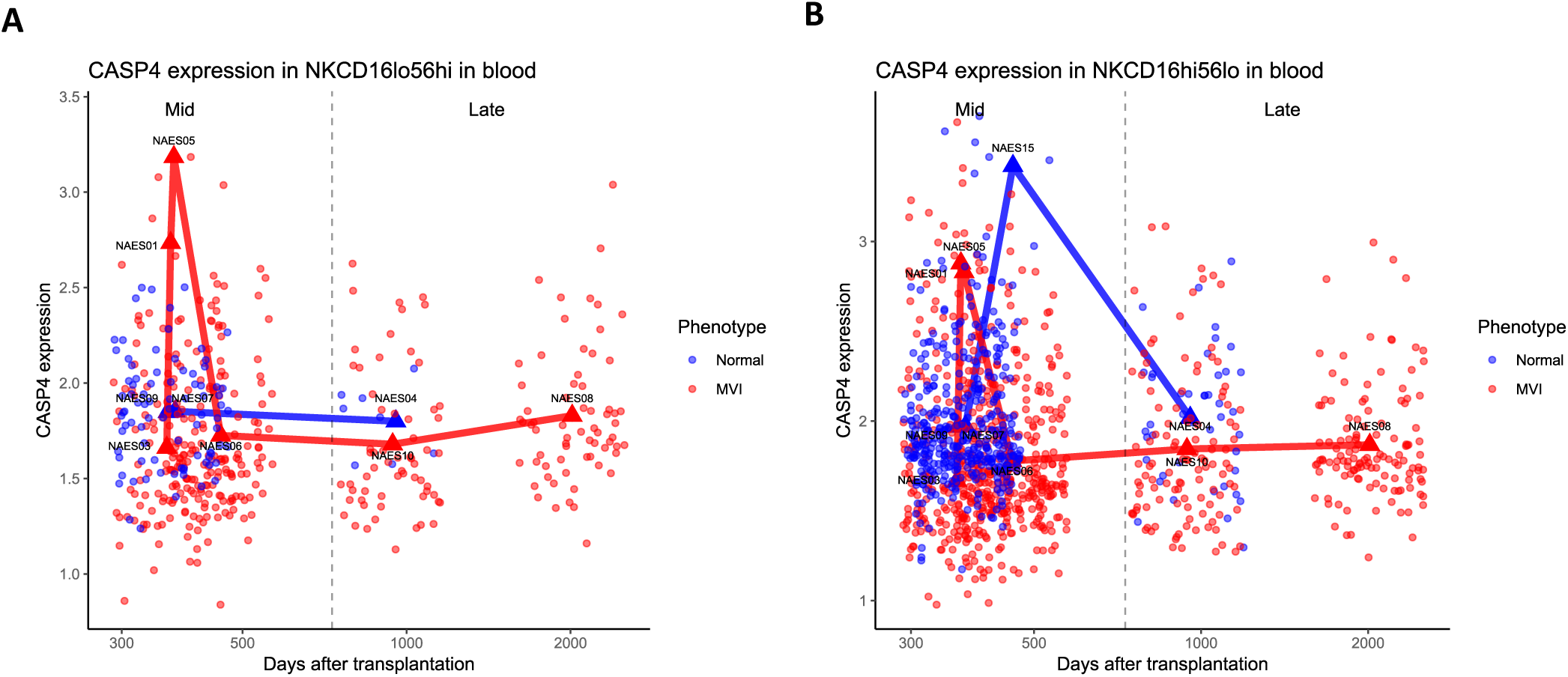
Caspase-4 expression in **A)** CD56^bright^CD16^low^ NK cells and **B)** CD56^dim^CD16^bright^ NK cells, according to time after transplantation in the different samples (*Mid*: 363-730 days; *Late*: >731 days).

## Notes

### Competing Interest Statement

The authors have declared no competing interest.

### Summary of Updates

We have included new experiments, new omics data and analysis of human samples from transplanted patients

