## Supplementary Tables for "Inflammasome Profiling in Human Natural Killer Cells under Pro-inflammatory Stimuli and Solid Organ Transplantation"

### Supplementary Table 1

| Patient #ID | Sex | Age (years) | Renal disease | Immune suppressive treatment | PRA | Donor Sex | Donor Age | Donor source |
|---|---|---|---|---|---|---|---|---|
| **#T1** | Male | 69 | CHK | Thymoglobulin+MMF++ST+Tacrolimus | 53% | Male | 55 | Deceased |
| **#T4** | Male | 73 | CHK | Thymoglobulin+MMF+ST+Tacrolimus | 0% | Male | 69 | Deceased |
| **#T5** | Male | 65 | CHK | Thymoglobulin+MMF+ST+Tacrolimuss | 0% | Male | 69 | Deceased |
| **#FJP** | Male | 47 | CHK | Thymoglobulin+MMF++ST+Tacrolimus | 0% | Female | 54 | Deceased |
| **#MFS** | Male | 68 | CHK | Baxiliximab+MMF+ST+Tacrolimus | 0% | Female | 54 | Deceased |
| **#ACS** | Male | 79 | CHK | Thymoglobulin+MMF+ST+Tacrolimus | 0% | Male | 65 | Deceased |
| **#MCB** | Female | 40 | CHK | Thymoglobulin+MMF+ST | 99% | Male | 65 | Deceased |

CHK: Chronic Kidney Disease; MMF: Mycophenolate Mofetil; ST: steroids; PRA: panel reactive antibodies.

### Supplementary Table 2

| Characteristic | Total (n=12) | MVI Group (n=6) | No MVI Group (n=6) | P-value | Test Used |
|---|---|---|---|---|---|
| Male sex n (%) | 8 (66.7) | 5 (83.3) | 3 (50.0) | 0.545 | Fisher's exact |
| Days after transplantation median (range) | 755 (364-2015) | 754 (364-2015) | 755 (366-2006) | 0.931 | Mann-Whitney U |
| Indication biopsy n (%) | 3 (25.0) | 3 (50.0) | 0 (0.0) | 0.182 | Fisher's exact |
| Glomerulitis (g) mean ± SD | 0.75 ± 0.62 | 1.5 ± 0.55 | 0.0 ± 0.0 | **0.001** | Mann-Whitney U |
| Interstitial inflammation (i) mean ± SD | 0.17 ± 0.39 | 0.33 ± 0.52 | 0.0 ± 0.0 | 0.250 | Mann-Whitney U |
| Tubulitis (t) mean ± SD | 0.42 ± 0.51 | 0.83 ± 0.41 | 0.0 ± 0.0 | **0.015** | Mann-Whitney U |
| Intimal arteritis (v) mean ± SD | 0.0 ± 0.0 | 0.0 ± 0.0 | 0.0 ± 0.0 | NA | NA |
| Peritubular capillaritis (cpt) mean ± SD | 0.83 ± 0.72 | 1.67 ± 0.52 | 0.0 ± 0.0 | **<0.001** | Mann-Whitney U |
| Transplant glomerulopathy (cg) mean ± SD | 0.67 ± 0.65 | 1.33 ± 0.52 | 0.0 ± 0.0 | **0.002** | Mann-Whitney U |
| Chronic interstitial fibrosis (ci) mean ± SD | 0.58 ± 0.51 | 1.0 ± 0.0 | 0.33 ± 0.52 | **0.041** | Mann-Whitney U |
| Chronic tubular atrophy (ct) mean ± SD | 0.75 ± 0.45 | 1.17 ± 0.41 | 0.33 ± 0.52 | **0.026** | Mann-Whitney U |
| Chronic vasculopathy (cv) mean ± SD | 0.83 ± 0.58 | 1.67 ± 0.52 | 0.5 ± 0.55 | **0.030** | Mann-Whitney U |
| Arteriolar hyalinosis (ah) mean ± SD | 1.08 ± 0.67 | 1.67 ± 0.52 | 0.5 ± 0.55 | **0.015** | Mann-Whitney U |
| Mesangial matrix increase (mm) mean ± SD | 0.25 ± 0.45 | 0.5 ± 0.55 | 0.0 ± 0.0 | 0.167 | Mann-Whitney U |
| Total inflammation (ti) mean ± SD | 0.42 ± 0.51 | 0.83 ± 0.41 | 0.0 ± 0.0 | **0.015** | Mann-Whitney U |
| C4d grade mean ± SD | 0.92 ± 0.90 | 1.83 ± 0.75 | 0.0 ± 0.0 | **0.004** | Mann-Whitney U |
| Total inflammation (ti) mean ± SD | 0.42 ± 0.51 | 0.83 ± 0.41 | 0.0 ± 0.0 | **0.015** | Mann-Whitney U |
| C4d grade mean ± SD | 0.92 ± 0.90 | 1.83 ± 0.75 | 0.0 ± 0.0 | **0.004** | Mann-Whitney U |

### Supplementary Table 3

| Variable | Total (n=9) | No MVI (n=5) | MVI (n=4) | p-value | Statistical Test |
|---|---|---|---|---|---|
| Demographics | | | | | |
| Sex (Male) n/total | 6/9 (66.7%) | 4/5 (80.0%) | 2/4 (50.0%) | 0.524 | Fisher's exact test |
| Time after transplantation (range in days) | 1155 (9-7709) | 1626 (35-7709) | 556 (9-2104) | 0.206 | Mann-Whitney U test |
| Clinical Characteristics | | | | | |
| Cause of biopsy (Indication) n/total | 7/9 (77.8%) | 4/5 (80.0%) | 3/4 (75.0%) | 1.000 | Fisher's exact test |
| Histological Scores (Mean) | | | | | |
| Acute lesions | | | | | |
| g (glomerulitis) | 0.56 | 0.00 | 1.25 | 0.016 | Mann-Whitney U test |
| i (interstitial inflammation) | 0.22 | 0.00 | 0.50 | 0.048 | Mann-Whitney U test |
| t (tubulitis) | 0.00 | 0.00 | 0.00 | 1.000 | Mann-Whitney U test |
| v (intimal arteritis) | 0.11 | 0.00 | 0.25 | 0.127 | Mann-Whitney U test |
| Chronic active lesions | | | | | |
| cpt (peritubular capillaritis) | 0.67 | 0.00 | 1.50 | 0.016 | Mann-Whitney U test |
| Chronic lesions | | | | | |
| cg (transplant glomerulopathy) | 0.00 | 0.00 | 0.00 | 1.000 | Mann-Whitney U test |
| ci (interstitial fibrosis) | 0.56 | 0.80 | 0.25 | 0.127 | Mann-Whitney U test |
| ct (tubular atrophy) | 0.67 | 0.80 | 0.50 | 0.413 | Mann-Whitney U test |
| cv (vascular fibrous intimal thickening) | 0.44 | 0.60 | 0.25 | 0.206 | Mann-Whitney U test |
| ah (arterial hyalinosis) | 0.22 | 0.40 | 0.00 | 0.048 | Mann-Whitney U test |
| Other lesions | | | | | |
| mm (mesangial matrix increase) | 0.00 | 0.00 | 0.00 | 1.000 | Mann-Whitney U test |
| ti (total inflammation) | 0.56 | 0.40 | 0.75 | 0.206 | Mann-Whitney U test |
| Immunostaining | | | | | |
| C4d grade | 0.22 | 0.00 | 0.50 | 0.048 | Mann-Whitney U test |
